# Conserved RNA helicase Vasa regulates ribonucleoprotein condensates through protein interaction and mRNA recruitment

**DOI:** 10.64898/2025.12.19.695503

**Authors:** Siran Tian, Hyosik Kim, Harrison A. Curnutte, Tatjana Trcek

## Abstract

Germlines contain ribonucleoprotein condensates known as germ granules, which concentrate proteins and mRNAs essential for animal development. Vasa, a conserved DEAD-box RNA helicase, is a core and highly concentrated constituent of germ granules. However, its roles within these structures remain poorly understood. Here, we use *Drosophila* germ granules as a model system to address this question. Applying in vivo and human cell systems, we found that condensation of Oskar (Osk) protein, which nucleates germ granules, occurred independently of Vasa. However, in oocytes lacking detectable Vasa protein, Osk formed aggregated condensates regardless of the eGFP tag. Furthermore, Osk-eGFP showed minimal recovery as measured by fluorescence recovery after photobleaching (FRAP) indicating that its exchange between condensates and their surroundings was greatly reduced in the absence of Vasa. Supporting this result, co-expression of Vasa increased the FRAP recovery of Osk-eGFP condensates and reduced Osk-eGFP partitioning into them in cells. This effect required the interaction between Vasa and Osk, suggesting that by binding Osk, Vasa modulates Osk’s phase behavior and its condensate material properties. Super-resolution microscopy further revealed that Vasa is required for germ granule mRNA localization to Osk condensates in vivo. Co-expression of Vasa with Osk-eGFP is necessary and sufficient to recruit germ granule mRNAs to condensates in cells. Although this activity depends on Vasa-Osk interaction, the interaction itself is not sufficient. Notably, localization of a conserved germ granule mRNA *nanos* reduced the FRAP recovery of Osk-eGFP condensates in cells, partially counteracting Vasa’s effect. Collectively, our findings uncovered a novel function of the DEAD-box RNA helicase Vasa in regulating the material properties of Osk condensates through coordinated protein-protein and protein-mRNA interactions.

**Four highlights:**

- Vasa modulates the material properties of Oskar ribonucleoprotein condensates in fly oocytes and human cells.
- Vasa modulates condensate material properties through its interaction with Oskar.
- mRNA localization to Oskar condensates requires Vasa and Vasa-Oskar interaction.
- *nanos* mRNA localization counteracts Vasa’s enhancement of Oskar exchange between condensates and their surroundings.

## Results and Discussion

Vasa is a conserved DEAD-box RNA helicase and serves as a canonical germline marker across diverse species.^1^ It localizes to ribonucleoprotein (RNP) condensates known as germ granules, which have been described across metazoans, including human,^2^ mouse,^3^ zebrafish,^4^ *C. elegans*,^5^ and *Drosophila*.^6^ Despite its evolutionary conservation and ubiquitous presence in the germline, Vasa’s function within germ granule RNP condensates remains unclear.

To investigate this, we used *Drosophila* germ granules, also called polar granules, as a model system. These RNP condensates form at the posterior of oocytes and embryos where they enrich specific proteins and mRNAs essential for development.^7,8^ Their formation is driven by the short isoform of Oskar (Short Osk), which nucleates germ granules, whereas the long isoform is excluded from germ granules and instead localizes to endocytic membranes to help anchor germ granules at the posterior.^9–11^ In addition, conserved RNA helicase Vasa, which interacts with Short Osk is also highly enriched in germ granules.^12–14^ Maternal loss of *vasa* expression prevents germ cell formation and triggers embryonic lethality.^6,15,16^ Despite being an established core granule component and promoter of fly development, the mechanisms underlying its importance in *Drosophila* germ granules have yet to be defined.

### *Drosophila* Vasa is dispensable for Osk condensation in vivo

DEAD-box RNA helicases, such as Vasa,^17,18^ have emerged as central regulators of ribonucleoprotein condensates, by modulating formation, material properties and intermolecular interactions among condensate components.^19–22^ Given that Osk nucleates and condenses germ granules and that Vasa physically interacts with its short isoform, we speculated that Vasa may influence germ granule assembly by modulating the condensation of Osk. To test this hypothesis, we examined whether loss of Vasa alters Osk condensation during germ granule formation. Specifically, we crossed flies expressing endogenous *oskar* (*osk*) gene tagged with eGFP (*osk-eGFP*) by CRISPR-Cas9 genome editing^23^ into a background lacking Vasa protein expression (*vasa^D1^*/*vasa^1^*)^24,25^ (Figure S1A). Although *osk-eGFP* marks both Osk isoforms, only the Short Osk condenses into germ granules.^9–11^ We analyzed oocytes expressing one copy of *osk-eGFP* and one copy of untagged *osk* (Figure S1A) following our previous quantitative analysis.^23^

Using anti-Oskar antibody staining to detect Osk in stage 10 oocytes, as well as in embryos before and after pole-cell formation (nuclear cycle (NC) 1–8 and 14, respectively), we confirmed that Osk-eGFP recapitulated the spatiotemporal localization of untagged endogenous Osk in *w¹¹¹⁸* flies (Figures 1A and S1B). Immunostaining could not be performed on stage 14 oocytes because the chorion blocks antibody access at this stage. Additionally, we and others have shown that the number and morphology of Osk-eGFP condensates closely resembled those of untagged Osk visualized by electron microscopy.^11,23,26,27^ Moreover, embryos carrying one copy of *osk-eGFP* and one untagged *osk* allele showed hatching rates comparable to *w¹¹¹⁸* controls.^23^ Thus, eGFP tagging does not markedly affect Osk localization, condensation or function.

**Figure 1.**
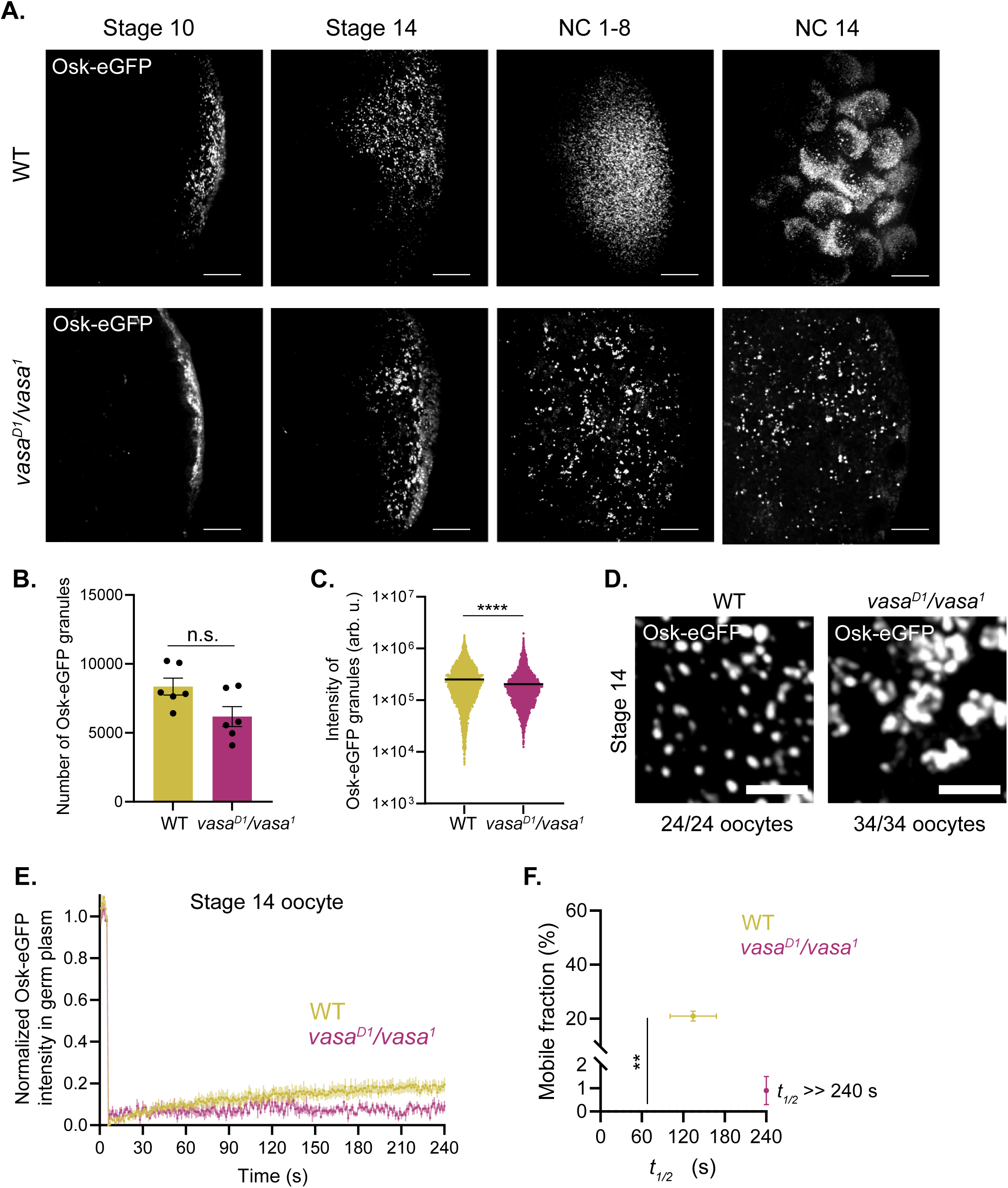
*Drosophila* Vasa modulates the material properties of Osk-eGFP condensates in late oocytes. (A) Accumulation of Osk-eGFP condensates at the posterior pole of oocytes (stages 10 and 14) and embryos (NC 1-8 and NC 14). Each image represents a maximal intensity projection from z-stacks. n = 8, 24, 15, and 16 oocytes or embryos for WT stage 10, stage 14, NC1-8, and NC 14, respectively. n = 15, 34, 16, and 6 for *vasa^D1^*/*vasa^1^*, respectively. Scale bars: 10 μm. (B) Average number of Osk-eGFP condensates per stage 14 oocyte. n = 6 oocytes for WT and *vasa^D1^*/*vasa^1^* oocytes. n.s.: not significant. p-value: 0.13 (two-tailed Mann-Whitney test). Data: mean ± SEM. (C) Distribution of Osk-eGFP condensate fluorescence intensity in stage 14 oocytes (n = 50128 and 28058 condensates analyzed for WT and *vasa^D1^*/*vasa^1^*, respectively). p-value (****): 3.28×10^−29^ (two-tailed t-test). Black line: average intensity. (D) Representative images of Osk-eGFP condensate morphology in stage 14 oocytes. Number of oocytes that demonstrate the shown phenotype is indicated. (E) Fluorescence recovery of Osk-eGFP in the germ plasm of stage 14 oocytes following photobleaching (n = 5 oocytes per genotype). Data: mean ± SEM from normalized fluorescence intensity values. (F) Analysis of mobile fraction (percentage of Osk-eGFP that exchanged between condensates and the surrounding environments) (%) and half-time recovery (*t_1/2_*; s) in germ plasm of WT and *vasa^D1^*/*vasa^1^* oocytes based on (E). p-value (**): 0.079 (mobile fraction; two-tailed Mann Whitney test). For *vasa^D1^*/*vasa^1^*, *t_1/2_* = 2.42×10^9^ ± 1.55×10^9^ s, which exceeds the experimental duration. Therefore, *t_1/2_* was set to 240 s in the figure. Because variability was high, *t_1/2_* values were not significantly different (p = 0.69; two-tailed Mann Whitney test). Data: mean ± SEM. See Methods.

Next, we examined the effect of Vasa expression on Osk condensation in vivo. Fluorescence microscopy showed that in ovaries derived from *vasa^D1^*/*vasa^1^* females, Osk-eGFP forms condensates independently of Vasa, from stage 10 oocytes through NC 14 embryos (Figure 1A), consistent with previous observations in *Drosophila* S2 cells.^26,28^ Western blot analysis confirmed that Vasa protein expression was not detectable in *vasa^D1^*/*vasa^1^* ovaries (Figures S1C and S1D), consistent with previous reports.^16,29^ Notably, Osk condensates also formed in oocytes and embryos from vasa*^D1^/*vasa^*1*^ flies carrying only untagged *osk* alleles (Figure S1E), indicating that condensation of Osk-eGFP was not driven by the eGFP itself. Therefore, our results revealed that Vasa is not required for Osk condensation in vivo.

### Vasa modulates the material properties of Osk-eGFP condensates in late oocytes

Next, we focused our analysis on stage 14 oocytes, the final stage of oogenesis. At this stage, cytoplasmic streaming has ceased^30^, and the germ plasm is functionally mature, with components such as Osk and Vasa stably incorporated into granules.^23,31^ This steady-state condition makes stage 14 suitable for quantitative analysis.

Using super-resolution microscopy coupled with a spot detection algorithm,^32^ we found that the number and intensity of Osk-eGFP condensates decreased by 26% and 19%, respectively in stage 14 oocytes derived from *vasa^D1^*/*vasa^1^* compared to wild-type (WT) controls (Figures 1B and 1C). This reduction agrees with the previous finding that loss of Vasa expression correlates with a reduced Short Osk abundance in ovaries (Figures S1C and S1D).^10^ Given that Osk condensate size approaches the resolution limit of our microscope (see Methods),^32,33^ the observed 19% decrease in fluorescence intensity may reflect a reduction in condensate volume, protein concentration or both. As these cannot be distinguished, we collectively refer to this as “protein partitioning into condensates” throughout the study.

Notably, in *vasa^D1^*/*vasa^1^* oocytes, Osk-eGFP condensates no longer presented as distinct puncta, but instead appeared aggregated, often connecting with each other into clusters of Osk condensates (Figure 1D). This phenotype persisted into NC 1-8 embryos (Figure S1Fi). Importantly, Osk condensates formed by untagged *osk* showed similar morphological changes in *vasa^D1^*/*vasa^1^* embryos (Figure S1Fii), indicating that aggregation is independent of the eGFP tag. Together, these results demonstrated that Vasa limits Osk condensate aggregation in vivo.

Because condensate morphology could reflect underlying material properties,^34^ we performed fluorescence recovery after photobleaching (FRAP) on stage 14 oocytes to measure Osk-eGFP exchange between condensates and their surroundings (Figure S1G), as described previously.^35^ We observed that in the presence of Vasa, 21.0 ± 1.8% of Osk-eGFP recovered by FRAP, with a half-time (*t_½_*) of 134.2 ± 33.4 seconds (s) (Figures 1E and 1F; see Methods). In the absence of Vasa, however, fluorescence recovery was nearly abolished, with only 0.9 ± 0.6% of Osk-eGFP exchanging with a *t_½_* exceeding the duration of the experiment (240 s) (Figure 1F). Notably, this loss of recovery was not observed in embryos laid by *vasa^D1^*/*vasa^1^* females (Figure S1H; Table S1), suggesting that in later developmental stages, embryo-specific factors regulate Osk-eGFP material properties independently of Vasa. Together, these results suggest that Vasa modulates the material properties of Osk-eGFP condensates in oocytes by reducing their aggregation and increasing their molecular exchange between condensates and their surroundings.

### Establishing a human cell line system to study the effect of Vasa on Osk condensates

Vasa performs several essential functions throughout oogenesis, including maintaining germline stem cells,^36,37^ stabilizing the piRNA processing complex,^25^ and regulating the Short Osk protein abundance,^10^ making it challenging to unambiguously dissect Vasa’s effect on Osk condensation that occurs at later stages of oogenesis. To circumvent this, we used human osteosarcoma (U2OS) cells, which lack *Drosophila* genes and the human Vasa homolog DDX4,^38^ providing a simplified cellular context to examine Vasa’s effect on Osk condensates.

We focused on Short Oskar, which condenses and recruits Vasa in flies^12–14^ and S2 cells.^26,28^ For expression, we used Short Osk that lacks the first nuclear localization signal (Short Osk ΔNLS1). In flies, this mutation increases cytoplasmic levels of Short Osk while minimally affecting germ granule assembly and egg hatching rate,^26^ indicating that Short Osk ΔNLS1 is functional. Using transient transfection, we co-expressed Short Osk ΔNLS1 tagged with eGFP (Short Osk ΔNLS1-eGFP) with Vasa tagged with a near-infrared fluorescent protein (Vasa-IRFP) (Figures 2Ai and 2Aii) in U2OS cells. Fluorescent tagging does not alter Vasa’s localization during oogenesis or embryogenesis,^23,39–41^ and flies expressing *vasa* tagged at its endogenous locus are homozygous viable,^40^ indicating largely intact activities of tagged Vasa.

**Figure 2.**
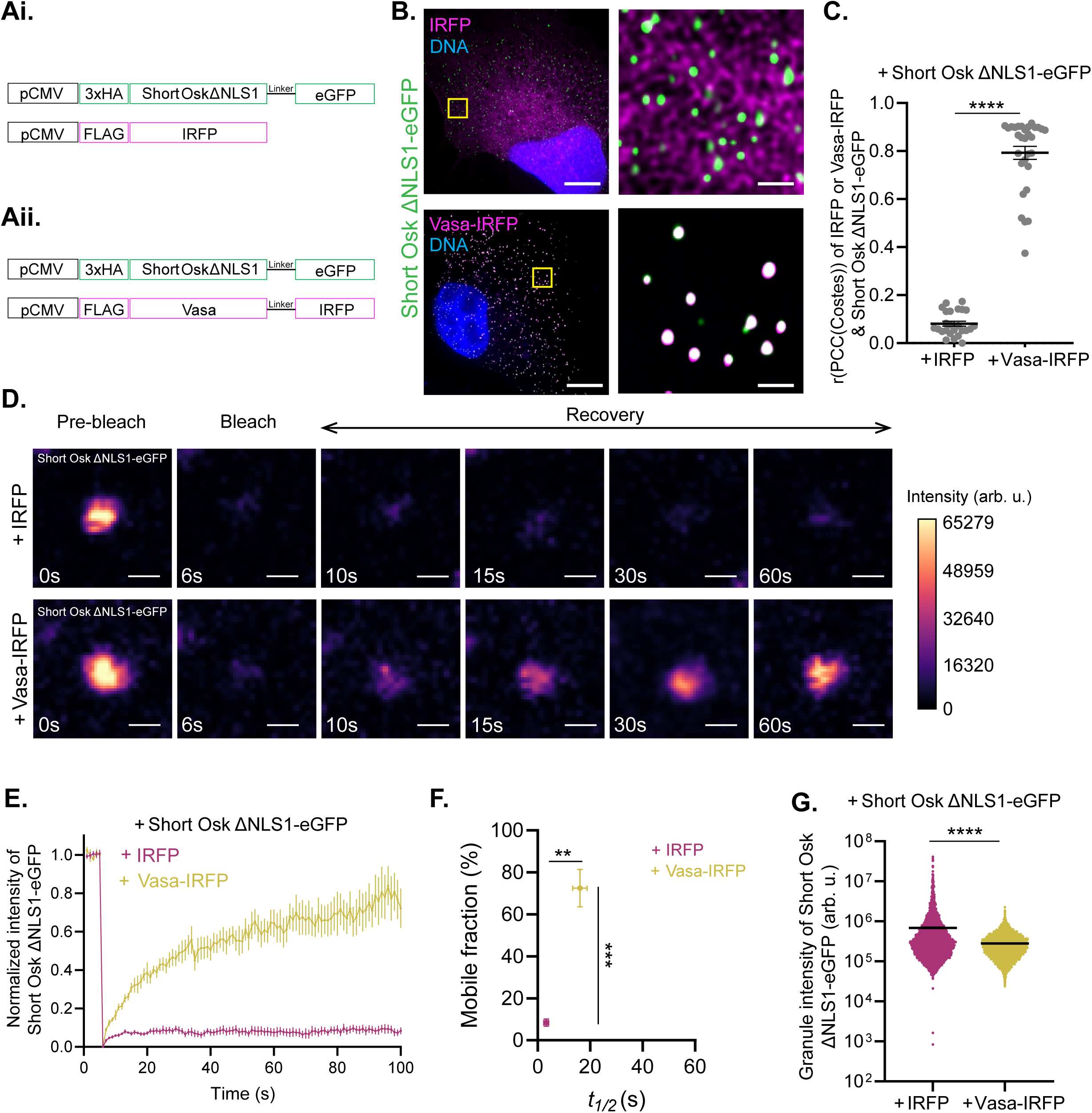
Vasa modulates the material properties of Short Osk ΔNLS1-eGFP condensates in cells. (A) Schematic of plasmid constructs used to co-express Short Osk ΔNLS1-eGFP with (i) IRFP or (ii) Vasa-IRFP. (B) Images of U2OS cells co-expressing Short Osk ΔNLS1-eGFP (green) with IRFP (top, magenta) or Vasa-IRFP (bottom, magenta). Yellow boxes: regions enlarged to the right. Nuclei were stained with DAPI (blue). Scale bars: 10 μm (left) and 1 μm (right). (C) Colocalization coefficients (r) determined using the PCC-Costes method. n = 25 and 30 cells for co-expression with IRFP and Vasa-IRFP, respectively. p-value (****): 4.71×10^−29^ (two-tailed t-test). Data: mean ± SEM. (D) Representative images of fluorescence recovery of Short Osk ΔNLS1-eGFP condensates in cells co-expressing IRFP (top) or Vasa-IRFP (bottom). Fluorescence intensities were normalized to the same scale. Scale bars: 0.5 μm. (E) Fluorescence recovery curves of Short Osk ΔNLS1-eGFP condensates after photobleaching. N = 7 and 8 condensates from different cells expressing IRFP or Vasa-IRFP, respectively. Data represent mean ± SEM of normalized fluorescence intensity values. (F) Mobile fraction (%) and *t_1/2_* (s) of Short Osk ΔNLS1-eGFP condensates based on (E). p-values: 0.0003 (mobile fraction; ***) and 0.0012 (*t_1/2_*; **) (two-tailed Mann Whitney test). Data: mean ± SEM. For IRFP, the SEM values for both mobile fraction and *t_1/2_* were too small to visualize. See Methods. (G) Quantification of Short Osk ΔNLS1-eGFP condensate fluorescence intensity in cells. n = 3778 and 8541 condensates analyzed for co-expression with IRFP and Vasa-IRFP, respectively. p-value (****): 1.11×10^−87^ (two-tailed t-test). Black line: average intensity.

Fluorescence microscopy showed that, as in vivo (Figure 1A) and in S2 cells,^26^ formation of Short Osk ΔNLS1-eGFP condensates occurred independently of Vasa in U2OS cells (Figure 2B). This condensation was not driven by the eGFP tag, as FLAG-tagged Short Osk ΔNLS1 also formed condensates without Vasa (Figure S2A). In contrast, without Osk, Vasa-IRFP did not condense (Figure S2B), consistent with Vasa-eGFP behavior in S2 cells.^26^

Unlike ovaries derived from *vasa^D1^/vasa^1^* females, which showed reduced Short Osk expression (Figures S1C and S1D), Short Osk ΔNLS1-eGFP levels were comparable in cells expressing Vasa-IRFP and IRFP alone (negative control) (Figures S2C-E). This suggested that the Vasa-dependent increase in Short Osk abundance in ovaries may require ovary-specific cofactors absent in U2OS cells, or that ΔNLS1 stabilizes Short Osk. Additionally, Vasa-IRFP colocalized with Short Osk ΔNLS1-eGFP, whereas IRFP did not (PCC(Costes): 0.79 ± 0.03 and 0.08 ± 0.01, respectively) (Figures 2B and 2C). Therefore, co-expression of Vasa-IRFP and Short Osk ΔNLS1-eGFP in U2OS cells reconstituted Osk condensates and recapitulated the spatial colocalization of Vasa and Short Osk observed in vivo and in S2 cells.^26,28,41^

### Vasa modulates the material properties of Short Osk ΔNLS1-eGFP condensates in cells

Next, we applied FRAP analysis to examine Short Osk ΔNLS1-eGFP condensates in U2OS cells. We observed that in the presence of Vasa-IRFP, Short Osk ΔNLS1-eGFP exhibited a mobile fraction of 72.5 ± 9.0% and a *t_½_* of 16.1 ± 2.8s (Figures 2E and 2F; Table S2). Notably, untagged Vasa generated a similar effect, suggesting that the increase in condensate mobile fraction is elicited by Vasa rather than the IRFP tag (Figures S2F and S2G; Table S3). In contrast, Short Osk ΔNLS1-eGFP co-expressed with IRFP exhibited a much lower mobile fraction (8.4 ± 1.8%) and a shorter *t_½_* (3.3 ± 0.9s) (Figures 2E and 2F; Table S2).

However, upon expression of Vasa-IRFP, both mobile fraction and *t_½_* in U2OS cells were higher and faster, respectively than those recorded in stage 14 oocytes (Figures 1E, 1F, 2E, and 2F). Moreover, when co-expressed with IRFP, Short Osk ΔNLS1-eGFP condensates appeared round and non-aggregated, unlike the irregular ones formed by Osk-eGFP in *vasa^D1^*/*vasa^1^* oocytes (Figures 1D and 2B). Furthermore, their fluorescence intensity was approximately 2.5-fold higher than in cells co-expressing Vasa-IRFP (Figure 2G), indicating a reduced partitioning of Short Osk ΔNLS1-eGFP into condensates upon Vasa-IRFP co-expression.

The morphological and FRAP recovery differences between late *vasa^D1^*/*vasa^1^* oocytes and cells expressing IRFP likely reflect distinct physiological contexts. While Osk condensate may recruit additional components to increase stability and preserve their morphology in late oocytes, U2OS cells may lack these factors. Alternatively, the NLS itself may help modulate the material properties of Osk condensates. Regardless of these differences, collectively, our data suggest that Vasa-IRFP modulates the material properties of Short Osk ΔNLS1-eGFP condensate by promoting molecular exchange with the surrounding environment while limiting protein accumulation in the condensate phase.

### Vasa modulates condensate material properties through its interaction with Short Osk

We next asked whether Vasa’s recruitment to Short Osk ΔNLS1-eGFP condensates is required for its ability to modulate the condensate material properties, or if its cytoplasmic presence alone is sufficient. To test this, we used previously characterized point mutations that disrupt the direct Vasa-Short Osk interaction in vitro. These included A162E and L228E in Short Osk and F504E and F508E in Vasa.^13,14^ Fluorescence microscopy confirmed that these mutations strongly reduced Vasa-IRFP localization to Short Osk ΔNLS1-eGFP condensates in U2OS cells (Figures 3A, 3B, S3A and S3B), consistent with the previous study done in S2 cells.^28^

**Figure 3.**
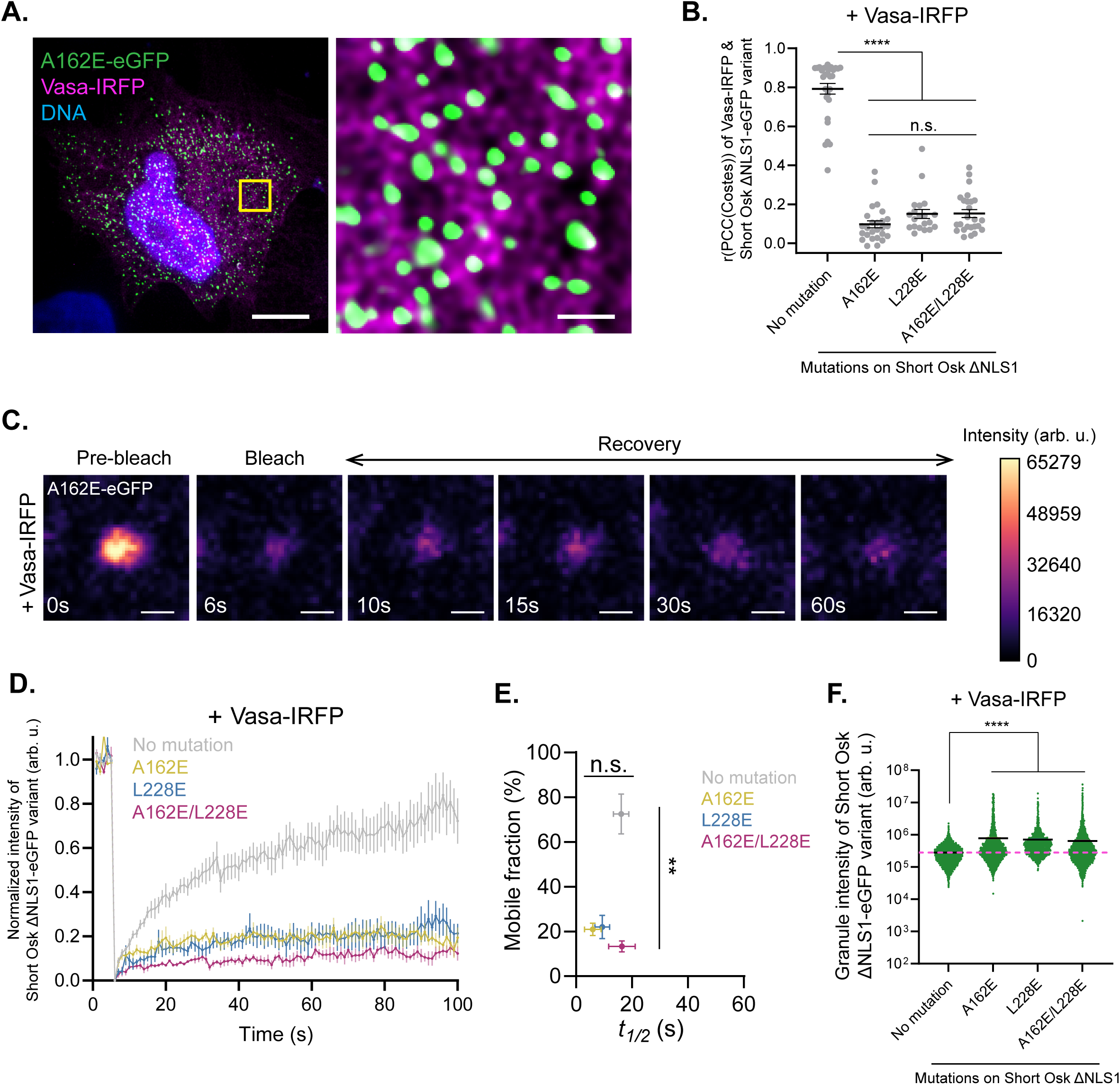
Vasa modulates condensate material properties through its interaction with Short Osk. (A) Images of U2OS cells co-expressing Short Osk ΔNLS1-eGFP carrying the A162E mutation (green) with Vasa-IRFP (magenta). Yellow box: region enlarged to the right. Nucleus was stained with DAPI (blue). Scale bars: 10 μm (left) and 1 μm (right). (B) Colocalization coefficients (r) determined using the PCC-Costes method. n = 30, 26, 20, and 25 cells expressing Short Osk ΔNLS1-eGFP with no additional mutation, A162E, L228E, and double mutations, respectively, together with Vasa-IRFP. n.s.: not significant. p-values (****; compared to no additional mutation (no mutation)): 2.57×10⁻²⁷ (A162E), 1.10×10⁻²¹ (L228E), and 1.56×10⁻²⁴ (A162E/L228E) (two-tailed t-test). p-value among A162E, L228E, and A162E/L228E: 0.08 (ANOVA test). Data: mean ± SEM. (C) Representative images of fluorescence recovery of the mutant Short Osk ΔNLS1-A162E-eGFP condensates in cells co-expressing Vasa-IRFP. Scale bars: 0.5 μm. (D) Fluorescence recovery of Short Osk ΔNLS1-eGFP condensates and its mutant variants co-expressed with Vasa–IRFP after photobleaching. n = 8, 6, 7, and 5 condensates from different cells expressing Short Osk ΔNLS1-eGFP with no mutation, A162E, L228E, or double mutations, respectively. Data represent mean ± SEM of normalized fluorescence intensity. (E) Mobile fraction (%) and *t_1/2_* (s) of Short Osk ΔNLS1-eGFP condensates based on (D). n.s.: not significant. For mobile fraction, p-values (vs. no mutation): 0.0013 (A162E; **), 0.0022 (L228E; **) and 0.0016 (A162E/L228E, **) (two-tailed Mann Whitney test). For *t_1/2_*, p-values (among all conditions): 0.062 (Kruskal-Wallis test). Data: mean ± SEM. See Methods. (F) Fluorescence intensity of Short Osk ΔNLS1-eGFP condensates and its mutant variants in cells co-expressing Vasa-IRFP. Data for the no mutation condition are from Figure 2G (+Vasa-IRFP, yellow). n = 8541, 2452, 1688, and 3761 condensates analyzed for Short Osk ΔNLS1-eGFP with no additional mutation, A162E, L228E, and double mutations, respectively, co-expressed with Vasa-IRFP. Magenta line indicates the average intensity of the no mutation condition. p-values (****; compared to no mutation condition): 8.52×10⁻¹²⁷ (A162E), 3.38×10⁻²¹⁰ (L228E), and 3.21×10⁻⁸¹ (A162E/L228E) (two-tailed t-test). Black line: average intensity.

FRAP analysis further revealed that co-expression of Vasa-IRFP with Short Osk ΔNLS1-eGFP carrying the A162E, L228E or A162/L228E mutations reduced the mobile fraction from 72.5 ± 9.0% to 21.0 ± 2.8%, 22.0 ± 5.3%, and 13.3 ± 2.4%, respectively (Figures 3C-3E; Table S2), with the corresponding *t_½_* values of 16.1 ± 2.8, 6.0 ± 3.0, 9.4 ± 2.6, and 16.4 ± 4.8s, respectively (Figure 3E; Table S2). Reciprocal experiments with Vasa-IRFP carrying the F504E, F508E, or F504E/F508E mutations produced similar results (Figures S3C and S3D; Table S2).

In these mutants, Short Osk ΔNLS1-eGFP condensates also exhibited higher intensity than those formed by co-expression of interaction-competent Osk and Vasa proteins, suggesting increased Short Osk partitioning into condensates (Figures 3F and S3E). These findings demonstrate that Vasa-Short Osk interaction is required for Vasa’s ability to enhance Short Osk ΔNLS1-eGFP exchange and decrease Short Osk partitioning into the condensate.

Finally, we examined Vasa point mutations known to impair its RNA-binding in vitro. These included O11 and O14, which impair RNA binding while retaining ATPase activity,^29^ and GNT, which abolishes both ATP and RNA binding.^24,25^ Consistent with observations in vivo,^24,29^ these mutations did not affect Vasa-IRFP localization to Short Osk ΔNLS1-eGFP condensates in U2OS cells (Figures S3F and S3G). FRAP recovery curves were also comparable between these Vasa mutants and WT Vasa-IRFP (Figures 2E, S3H and S3I; Table S2). Moreover, condensate fluorescence intensities were similar to or lower than those with WT Vasa-IRFP (Figure S3J). Thus, mutations that disrupt Vasa RNA- and ATP-binding in vitro do not impair Vasa’s ability to modulate the material properties of Short Osk ΔNLS1-eGFP condensates in cells.

### Vasa is critical for germ granule mRNA localization to Osk condensates in vivo

In *Drosophila*, mRNAs such as *nanos* (*nos*), *polar granule component* (*pgc*), and *germ cell-less* (*gcl*) localize to germ granules formed by Osk and are required for embryonic patterning and germ cell formation.^41–46^ Since maternal loss of *vasa* expression disrupts embryonic patterning and abolishes germ cell formation,^6,15,16^ we speculated that these defects could be due to impaired mRNA localization.

Using single-molecule fluorescent in situ hybridization (smFISH), we examined *nos*, *pgc*, and *gcl* mRNA localization to Osk condensates in WT and *vasa^D1^/vasa^1^* stage 10 to stage 14 oocytes (Figure S1A). In *vasa^D1^/vasa^1^*, only 60% of stage 10 oocytes exhibited *nos* localization, and all lost detectable posterior localization by stage 14 oocytes (Figures 4A, S4Ai and S4Aii). In contrast, WT oocytes robustly localized *nos* to germ granules beginning at stage 10 of oogenesis (Figure 4A), consistent with previous observations.^23,45^ Similar trends were also observed for *pgc* and *gcl* (Figures S4Ai and S4Aii). Importantly, Osk-eGFP condensates persisted through oogenesis (Figure 1A), indicating that loss of mRNA localization was not due to lack of Osk condensation. Together, these results demonstrate that Vasa is required for robust and sustained mRNA localization to Osk condensates in vivo.

**Figure 4.**
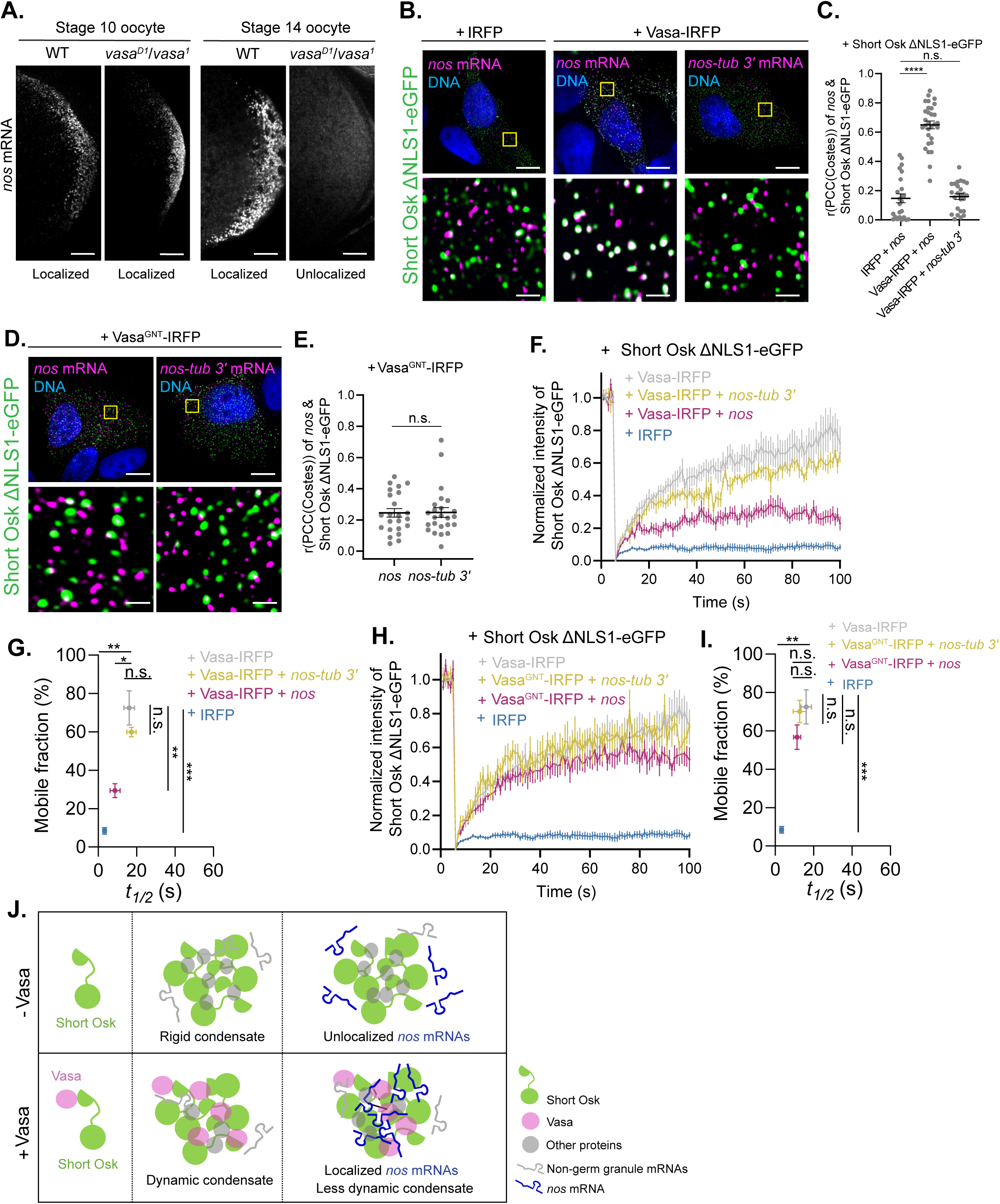
*nos* mRNA localization counteracts Vasa’s enhancement of Short Osk ΔNLS1-eGFP exchange between condensates and their surroundings. (A) smFISH images of *nos* mRNA at the posterior of stage 10 and stage 14 oocytes in WT and *vasa^D1^*/*vasa^1^*. Each image represents a maximal intensity projection from z-stacks. Scale bars: 10 μm. (B) smFISH images of *nos* mRNA (magenta) in cells co-expressing Short Osk ΔNLS1-eGFP with IRFP (left) or Vasa-IRFP (middle), and of *nos-tub* 3′ mRNA (magenta) in cells co-expressing Short Osk ΔNLS1-eGFP (green) and Vasa-IRFP (right). Yellow boxes: regions enlarged at the bottom. Nuclei were stained with DAPI (blue). Scale bars: 10 μm (top) and 1 μm (bottom). (C) Colocalization coefficients (r) determined using the PCC-Costes method. n = 24, 30, and 25 cells c0-expressing Short Osk ΔNLS1-eGFP with IRFP and *nos*, Vasa-IRFP and *nos*, and Vasa-IRFP and *nos–tub* 3′, respectively. n.s.: not significant. p-values (compared to IRFP + *nos*): 3.18×10⁻¹⁷ (****; Vasa-IRFP + *nos*) and 0.74 (Vasa-IRFP + *nos-tub* 3′) (two-tailed t-test). (D) smFISH images of *nos* and *nos-tub* 3′ mRNAs (magenta) in cells co-expressing Short Osk ΔNLS1-eGFP (green) and Vasa-IRFP, which carries the GNT mutation. Yellow boxes: regions enlarged at the bottom. Nucleus were stained with DAPI (blue). Scale bars: 10 μm (top) and 1 μm (bottom). (E) Colocalization coefficients (r) determined using the PCC-Costes method. n = 23 and 24 cells co-expressing Short Osk ΔNLS1-eGFP and Vasa-IRFP carrying the GNT mutation, together with *nos* and *nos-tub* 3′ mRNAs, respectively. n.s.: not significant. p-value: 0.94 (two-tailed t-test). Data: mean ± SEM. (F) Fluorescence recovery of Short Osk ΔNLS1-eGFP condensates after photobleaching. Data for Vasa-IRFP and IRFP are from Figure 2E. Mean ± SEM of normalized fluorescence intensity values from n = 7, 7, 7, and 8 condensates from different cells expressing Vasa-IRFP, Vasa-IRFP and *nos-tub* 3′, Vasa-IRFP and *nos*, or IRFP together with Short Osk ΔNLS1-eGFP is shown, respectively. (G) Mobile fraction (%) and *t_1/2_* (s) of Short Osk ΔNLS1-eGFP condensates derived from (F). n.s.: not significant. For mobile fraction, p-values (vs. Vasa-IRFP): 0.18 (Vasa-IRFP + *nos-tub* 3′), 0.0059 (Vasa-IRFP + *nos*; **) and 0.0003 (IRFP; ***) (two-tailed Mann Whitney test). For *t_1/2_*, p-values (vs. Vasa-IRFP): 0.69 (Vasa-IRFP + *nos-tub* 3′), 0.040 (Vasa-IRFP + *nos*; *) and 0.0012 (IRFP; **) (two-tailed Mann Whitney test). Data: mean ± SEM. For IRFP, the SEM values for both mobile fraction and *t_1/2_* were too small to visualize. See Methods. (H) Fluorescence recovery of Short Osk ΔNLS1-eGFP condensates after photobleaching. Data for Vasa-IRFP and IRFP are the same as in Figures 2E and 4F. Mean ± SEM of normalized fluorescence intensity values of n = 7, 6, 7, and 8 condensates from different cells expressing Vasa-IRFP, Vasa^GNT^-IRFP and *nos-tub* 3′, Vasa^GNT^-IRFP and *nos*, or IRFP together with Short Osk ΔNLS1-eGFP is shown, respectively. (I) Mobile fraction (%) and *t_1/2_* (s) of Short Osk ΔNLS1-eGFP condensates derived from (F). n.s.: not significant. For mobile fraction, p-values (vs. Vasa-IRFP): 0.92 (Vasa^GNT^-IRFP + *nos-tub* 3′), 0.27 (Vasa^GNT^-IRFP + *nos*) and 0.0003 (IRFP; ***) (two-tailed Mann Whitney test). For *t_1/2_*, p-values (vs. Vasa-IRFP): 0.57 (Vasa-IRFP + *nos-tub* 3′), 0.40 (Vasa-IRFP + *nos*) and 0.0012 (IRFP; **) (two-tailed Mann Whitney test). Data: mean ± SEM. For IRFP, the SEM values for both mobile fraction and *t_1/2_* were too small to visualize. See Methods. (J) Proposed model illustrating how Vasa may regulate Short Osk condensation. In the absence of Vasa, Short Osk (green) condenses, perhaps with other components (grey), but these condensates are rigid and fail to recruit germ granule mRNAs, such as *nos* (blue). In contrast, Vasa (pink) renders the condensates more dynamic and enables recruitment of *nos* mRNAs. However, Osk condensates containing localized *nos* mRNAs are less dynamic. Therefore, *nos* mRNA localization partially counteracts Vasa’s effects on condensates.

### Vasa and Short Osk are sufficient for localizing germ granule mRNAs in cells

We next asked whether Vasa and Short Osk are sufficient to localize germ granule mRNAs. To test this, we co-expressed Short Osk ΔNLS1-eGFP with either IRFP or Vasa-IRFP together with *nos*, *pgc*, or *gcl* in U2OS cells. Using smFISH coupled with super-resolution microscopy, we quantified colocalization between these mRNAs and Short Osk ΔNLS1-eGFP. In IRFP-expressing cells, colocalization coefficients for *nos*, *pgc*, and *gcl* with Short Osk ΔNLS1-eGFP condensates were 0.15 ± 0.03, 0.07 ± 0.02, and 0.18 ± 0.02, respectively (Figures 4B, 4C, S4B and S4C). We observed that co-expression with Vasa-IRFP markedly increased these values to 0.65 ± 0.03, 0.50 ± 0.03, and 0.47 ± 0.04, respectively (Figures 4B, 4C, S4D and S4E), independent of mRNA expression levels (Figures S4F). Notably, the colocalization pattern mirrored that recorded in WT embryos, with *nos* showing the highest colocalization, followed by *pgc* and *gcl.*^41^ Therefore, Vasa and Short Osk are sufficient not only to promote mRNA recruitment to condensates but also recreate the spatial patterning hierarchy observed in embryos within U2OS cells.

Because the *nos* 3′UTR is required for *nos* localization in embryos,^47^ we replaced it with the 3′UTR of *bicoid* (*bcd*) or *tubulin* (*tub*), both of which abolish posterior *nos* localization in embryos^42,47^. As expected, *nos-bcd* 3′UTR and *nos-tub* 3′UTR mRNAs greatly reduced colocalization with Short Osk ΔNLS1-eGFP when co-expressed with Vasa-IRFP (PCC(Costes): 0.13 ± 0.03 and 0.16 ± 0.02, respectively) (Figure S4E), revealing that co-expression of Short Osk ΔNLS1-eGFP and Vasa-IRFP in U2OS cells recapitulated the mRNA localization specificity observed in embryos.^48^ Together, these results show that co-expression of Vasa-IRFP and Short Osk ΔNLS1-eGFP is sufficient for germ granule mRNA localization to condensates in U2OS cells.

### Vasa-Short Osk interaction is required but not sufficient for robust *nos* localization in cells

To determine whether mRNA localization depended on Vasa-Short Osk interaction or Vasa’s cytoplasmic presence itself, we focused on *nos* mRNA, which showed the strongest colocalization with Short Osk ΔNLS1-eGFP (Figure S4E). This provided a sensitive readout for detecting localization changes caused by mutations in either Short Osk or Vasa.

smFISH and super-resolution microscopy revealed that disrupting the Vasa-Short Osk interaction via point mutations (Short Osk: A162E, L228E and A162E/L228E; Vasa: F504E, F508E and F504E/F508E)^13,14^ significantly reduced *nos* mRNA localization to levels comparable to that observed in the IRFP control (dashed purple line) (Figures S4G-S4I). Therefore, the Vasa-Short Osk interaction is required for *nos* localization to condensates.

In contrast, co-expression of Short Osk ΔNLS1-eGFP with Vasa-IRFP carrying the O11, O14 or O11/O14 mutations decreased *nos* colocalization to intermediate levels between IRFP and Vasa-IRFP (Figures S4J and S4K). Notably, *nos* localization was severely impaired in cells expressing Vasa^GNT^-IRFP (PCC(Costes): 0.24 ± 0.03), reaching a level comparable to the *nos-tub* 3′UTR control (PCC(Costes): 0.25 ± 0.03) (Figures 4D and 4E). Although O11, O14, and GNT mutations did not affect Vasa interaction with Short Osk ΔNLS1-eGFP and its recruitment into condensates (Figures S3F and S3G), GNT caused the most severe *nos* localization defect. These results indicated that while the Vasa-Short Osk interaction is necessary for *nos* mRNA localization to condensates, it is not sufficient. Therefore, additional Vasa activities are essential for robust *nos* mRNA localization to Short Osk ΔNLS1-eGFP condensates in cells.

### *nos* mRNA localization counteracts Vasa’s enhancement of Short Osk ΔNLS1-eGFP exchange

Given Vasa’s roles in increasing the mobile fraction of Short Osk ΔNLS1-eGFP condensates and promoting mRNA localization to them in U2OS cells (Figures 2E and 4B), we next asked whether mRNA localization in turn influences condensate FRAP recovery. To do this, we generated a construct that co-expressed *nos* and Short Osk ΔNLS1-eGFP.

FRAP analysis showed that, when co-expressed with Vasa-IRFP, Short Osk ΔNLS1-eGFP condensate had a mobile fraction of 72.5 ± 9.0% and a *t_½_* of 16.1 ± 2.8s (Figure 2F). However, upon additional *nos* expression, the mobile fraction dropped to 29.4 ± 3.6% while *t_½_* decreased to 8.7 ± 2.6s (Figures 4F and 4G; Table S2). Nevertheless, the condensates still recovered more than in cells co-expressing only IRFP (Figures 4F and 4G). For comparison, co-expression with *nos-tub* 3′UTR, which poorly localized to Short Osk ΔNLS1-eGFP condensates (Figures 4B and 4C), exhibited a mobile fraction of 59.9 ± 2.5% and a *t_½_* of 17.3 ± 2.5s, similar to those recorded in the absence of *nos* mRNA expression (Figures 4F and 4G; Table S2).

To determine whether the FRAP changes were due to *nos* localization or its cytoplasmic presence, we examined condensates in cells expressing *nos* and Vasa^GNT^-IRFP. The GNT mutation retained Vasa’s ability to modulate condensates (Figures S3H and S3J), but disrupted *nos* localization (Figures 4D and 4E). These condensates exhibited a mobile fraction of 56.7 ± 6.3% and a *t_½_* of 11.4 ± 1.8 s, comparable to those recorded in cells that did not express *nos* mRNA (Figures 4H and 4I; Table S2).

Together, these data revealed that localized *nos* mRNA counteracts Vasa’s effects on condensates by partially reducing the exchange of Short Osk ΔNLS1-eGFP between condensates and their surroundings. However, in the presence of both *nos* and Vasa-IRFP, Short Osk ΔNLS1-eGFP condensates still remained relatively dynamic, in contrast to the rigid, near-immobile condensates observed in cells that only co-expressed IRFP (Figure 4J).

## Conclusion

In this study, we investigated how the conserved RNA helicase Vasa regulates *Drosophila* Osk condensates. We observed that in late oocytes, Vasa enhances Osk exchange between condensates and their surroundings, and prevents their aggregation (Figures 1D and 1E). In human cells, Vasa not only enhances Osk exchange but also reduces the amount of Osk partitioned into the condensate phase (Figures 2E and 2G), effects that require Vasa’s interaction with Osk. Furthermore, Vasa is critical for germ granule mRNA localization to Osk condensates in vivo, and co-expression of Vasa with Osk is sufficient to drive this localization in cells (Figures 4A-C). Notably, *nos* localization counteracts Vasa’s enhancement of condensate recovery (Figure 4F), consistent with previous observations that RNAs modulate condensate material properties.^49,50^ Therefore, Vasa regulates the material properties of Osk condensates through coordinated protein-protein and protein-RNA interactions.

Furthermore, mutations in Vasa that disrupt RNA binding in vitro (O11, O14, and GNT) do not significantly alter its interaction with Osk in cells (Figures S3F and S3G), consistent with previous in vivo observations.^24,29^ We also observe that *nos* mRNA localization is only moderately affected by the O11 and O14 mutants (Figures S4J and S4K), while the GNT mutant causes a stronger defect (Figures 4D and 4E). However, this does not imply that mRNA localization is independent of Vasa’s RNA-binding activity. The difference between in vitro and in cellulo results likely arose from cofactors such as Osk that can interact with Vasa and which are present in cells and absent in vitro. Such interactions could rescue the loss of Vasa’s RNA-binding activity caused by point mutations, as proposed previously.^51^ Alternatively, germ granule mRNAs may primarily associate with Short Osk, with Vasa modulating Short Osk conformation to facilitate mRNA recruitment. Further studies are required to define the mechanisms of mRNA recruitment by Osk condensates.

DEAD-box RNA helicases maintain condensate integrity by remodeling intramolecular and intermolecular RNA-RNA interactions.^19–22^ Beyond this canonical role, our results reveal that Vasa directly shapes condensate properties through its interplay with both proteins and RNAs. By orchestrating intermolecular protein-protein, protein-RNA, and RNA-RNA interactions, Vasa serves as a multifaceted regulator that controls condensate biophysical properties and composition and safeguards against aberrant or pathological condensate formation. Given the evolutionary conservation of Vasa, this work offers broader insights into the functions of DEAD-box RNA helicases across germ granules.

## METHODS

### EXPERIMENTAL MODEL AND SUBJECT DETAILS

#### Fly stocks

All fly stocks and crosses were raised on standard cornmeal agar media at 25℃. To quantify the number and fluorescence intensity of short Osk-eGFP granules, we used eggs laid by females expressing Osk genetically modified with CRISPR/Cas9 at the endogenous locus with eGFP. We used heterozygous *osk-eGFP* for our experiments following our previous quantitative analysis.^23^ To generate flies lacking Vasa proteins, we used two *vasa* alleles *vasa^D1^* and *vasa^1^*. While *vasa^1^* produces Vasa protein detectable in the germarium of adult ovaries, *vasa^D1^* has no detectable Vasa protein in ovaries.^16^ Both alleles encode identical amino acid sequences to WT Vasa.^29^ Females carrying *vasa^1^/cyo; osk-eGFP/TM3sb* (*vasa^1^/cyo*; BDSC: 1574) were crossed with males expressing *vasa^D1^/cyo; osk-eGFP/TM3sb* (*vasa^D1^/cyo*; gift from Ruth Lehmann lab). The resulting *vasa^1^/vasa^D1^; osk-eGFP/TM3sb* progeny were used for experiments.

#### Cell culture

U2OS cells (ATCC: HTB-96) were cultured in Dulbecco’s Modified Eagle Medium (DMEM) (Corning: 10-013-CV) supplemented with 10% Fetal Bovine Serum (FBS) (Corning: 35-011-CV) and 1X penicillin-streptomycin (Thermo Fisher: 15070063). Cells were maintained at 37°C in a humidified incubator with 5% CO_2_.

### METHODS DETAILS

#### Microscopes

Both fixed fly samples and cells were imaged in 3D with a z-step size of 150 nm for colocalization analysis and quantification of condensate number or intensity. Imaging was performed using a HC PL APO 100x/1.47 OIL CORR TIRF oil objective on a vt-instant Structured Illumination Microscope (vt-iSIM; BioVision Technologies). 405 nm, 488 nm, 561 nm and 642 nm lasers were used.

For FRAP assays, live-cell imaging was performed using a Zeiss LSM980 laser scanning confocal microscope equipped with a temperature-controlled chamber. Oocyte and embryo FRAP experiments were done at room temperature, whereas FRAP in cultured cells was performed at 37℃. Images were acquired with a PL APO 63×/1.4 DIC oil objective using a 488 nm laser.

#### Immunostaining of fly ovaries

Females were fed with yeast powder the day before the dissection. Ovaries were dissected at room temperature (RT) in Schneider’s insect medium supplied with 10% FBS, 1X penicillin-streptomycin and 200 μg/mL insulin (Sigma: I5500-500MG).^23^ Samples were fixed in 4% paraformaldehyde (Electron Microscopy Sciences: 15713) in 0.1% PBTx (1X PBS, 0.1% Triton-X100 (Millipore: TX1568-1) for 20 minutes (mins) at RT, followed by two 10-min washes in 0.1% PBTx.^52^

Afterwards tissues were blocked for 30 mins at RT in blocking buffer (0.1% PBTx, 0.5% BSA (Millipore: A3294-50G), 2% normal goat serum (Abcam: ab7481)). Primary antibody incubation was carried out overnight at 4℃ using rabbit anti-Osk antibody (gift from Anne Ephrussi lab) diluted 1:500 in the blocking buffer. Afterward, samples were washed five times in 0.1% PBTx for 10 mins each at RT and re-blocked in the blocking buffer for 30 mins at RT.

Incubation with the secondary antibody lasted 4 hours (hrs) at RT using goat anti-rabbit IgG Alexa 488 antibody (Thermo Fisher: A-11070) diluted 1:1000 in the blocking buffer. Samples were washed five times in 0.1% PBTx for 10 mins each. DNA was stained with 1 μg/mL DAPI (Sigma: 10236276001) diluted in 0.1% PBTx for 10 mins at RT. Finally, samples were mounted on slides using ProLong™ glass antifade mountant (Thermo Fisher: P36980) and cured overnight at RT before imaging.

#### Immunostaining of fly embryos

Embryos were collected as described before^32^ and stored in 100% methanol (Thermo Fisher: A412-4) at 4℃. To rehydrate embryos, they were washed twice in 50% methanol diluted in 0.3% PBTx (1X PBS, 0.3% Triton-X100) for 5 mins at RT. Then embryos were washed and permeabilized in 0.3% PBTx for 15mins, repeated three times at RT.

The subsequent immunostaining procedure followed the same steps as described for ovaries (see above), except that the blocking buffer contained 0.3% PBTx, 1% BSA and 2% normal goat serum. All washes were performed using 0.3% PBTx.

#### Western blotting in fly ovaries

Twenty-five to thirty well-fed females aged three to seven days were anesthetized with CO_2_ and dissected in Schneider’s insect medium supplemented with 10% FBS, 1X penicillin-streptomycin and 200 μg/mL insulin (Sigma: I5500-500MG).^23^ Ovaries were rinsed quickly in 1X PBS, flash-frozen in liquid nitrogen and stored in −80℃ until use. For lysis, ovaries were homogenized in 100 μL of cold lysis buffer (150 mM sodium chloride (Thermo Fisher: S271-500), 50 mM pH 8.0 Tris-HCl (Promega: H5121), 1% Triton-X100, 0.5% sodium deoxycholate (Sigma: 30970-25G), and 0.1% SDS (Thermo Fisher: 15553027)). Lysates were then incubated on ice for 20 mins. Downstream procedures for western blotting were performed as described previously.^52^ For fly experiments, the following antibodies were used: primary antibodies rabbit anti-Vasa (gift from Ruth Lehmann Lab; 1:4000), rabbit anti-Osk (gift from Anne Ephrussi Lab; 1:5000), mouse anti-α-tubulin (DSHB: 12G10; 1:1000), and secondary antibodies goat anti-rabbit IgG (HRP) (Abcam: ab6721; 1:10000) and goat anti-mouse IgG (HRP) (Abcam: ab6789; 1:10000).

#### Fluorescence recovery after photobleaching (FRAP) of live oocytes and embryos

For live imaging of oocytes, about 10 well-fed females aged three to seven days were anesthetized with CO_2_ and dissected in Schneider’s insect medium supplemented with 10% FBS and 1X penicillin-streptomycin. Dissection forceps were used to tease apart the connective tissue holding the ovarioles together. Ovaries were glued to a coverslip and mounted in a Lumox imaging chamber (Sarstedt: 94.150.101) and embedded with Halocarbon 200 oil (Halocarbon: 200). For embryo imaging, embryos were collected on apple juice-agar plates after 1 hr of egg laying at 25 °C. They were dechorionated in 50% bleach for 1-2 minutes, rinsed through a Nitex nylon mesh, and washed twice with water. Embryos were then glued to a coverslip using the same procedure as for ovaries.

Because individual germ granules were smaller than the resolution limit of the microscope and therefore difficult to examine with FRAP individually, a 3 μm × 3 μm region of interest (ROI) containing germ plasm and germ granules was studied instead, as done previously.^26,35^ To establish the pre-bleach state, samples were imaged for 5 seconds at 1-second intervals and then bleached with a strong 488-nm laser pulse. Recovery was recorded by acquiring images every second thereafter as described previously.^35^

#### Transfection of cells

Xfect transfection reagent (Takara: 631318) was used following the manufacturer’s instruction. Specifically, one day before transfection, 1.2×10^5^ cells were seeded either into a well of a 6-well plate (Corning: 351146) or into a 30 mm glass bottom dish (CELLTREAT: 229632). If the cells were used for smFISH experiments, each well also contained a collagen-coated coverslip (Neuvitro: GG-18-1.5-COLLAGEN). On the day of transfection, a total of 5 μg DNA (containing one or two plasmids) or 7.5 μg DNA (for co-transfection with three plasmids) was mixed with Xfect reagent and added to the cells. After 4-5 hrs, the medium was replaced with fresh DMEM supplemented with 10% FBS and 1X penicillin-streptomycin. Cells were allowed to recover for 24 hrs before fixation, sample collection for western blotting or live-cell imaging.

#### Immunostaining of cells

Cells were seeded on collagen-coated coverslips as described above and washed three times with 1X PBS at RT. Samples were fixed in 4% paraformaldehyde for 10 mins at RT followed by three washes in 1X PBS for 5 mins each. Cells were then permeabilized in 0.1% PBTx for 5 mins at RT and blocked for 30 mins in blocking buffer containing 1% BSA diluted in 0.1% PBTx.

Primary antibody incubation was performed overnight at 4℃ using rabbit anti-FLAG (Abcam: AB205606) diluted 1:500 in the blocking buffer. Samples were washed three times with 1X PBS for 10 mins each, followed by secondary antibody incubation for 2 hrs at RT using goat anti-rabbit IgG Alexa 647 (Thermo Fisher: A32733) diluted 1:1000 in the blocking buffer. Samples were washed three times with 1X PBS for 10 mins each. DNA was stained with 1 μg/mL DAPI diluted in 1X PBS for 10 mins at RT. Finally, samples were mounted on slides using ProLong™ glass antifade mountant and cured overnight at RT before imaging.

#### Western blotting in cells

Cells growing in a 6-well plate were quickly washed with 1ml of 1X PBS. Afterwards, 1 mL 1X PBS was added to each well. Cells were then scraped with a cell scraper and transferred to 1.5 mL tubes. Samples were centrifuged at 15,000 g for 3 minutes to remove PBS, and the resulting pellets were flash-frozen in liquid nitrogen and stored at −80 °C until use. Pellets were resuspended in 50 μL of lysis buffer (150 mM sodium chloride, 50 mM Tris (pH 7.0) (Millipore: 648314-100ML), 1% Triton-X100) and incubated on a rotator for 20 mins at 4℃. Cell lysates were centrifuged at 15,000 g for 10 mins at 4℃ to remove the cellular debris. Downstream procedures for western blotting were performed as described previously.^52^ For cell experiments, the following antibodies were used: primary antibodies rabbit anti-FLAG (Abcam: ab205606; 1:5000), mouse anti-HA (Thermo Fisher: 26183; 1:5000) and mouse anti-β-Actin (Abcam: ab8224; 1:5000), and secondary antibodies goat anti-rabbit IgG (HRP) (1:10000) and goat anti-mouse IgG (HRP) (1:10000).

#### FRAP of live cells

As described in Methods - Transfection of cells, after 24 hours of transfection on a 30 mm glass-bottom dish, cells were washed twice with 1× PBS. 1 mL of Live Cell Imaging Solution (Thermo Fisher: A59688DJ) supplemented with 10 μL of Prolong Live Antifade Reagent (Thermo Fisher: P36975) was added, and cells were incubated at 37℃ for 30 mins.

For FRAP of short Osk ΔNLS1-eGFP granules in cells, an ROI was drawn as tightly as possible around the granule of interest. Imaging was then performed as described above.

#### smFISH in fly oocytes

Fifteen to twenty-five well-fed females aged three to seven days were anesthetized with CO_2_ and dissected in Schneider’s insect medium (Sigma: S0146) supplemented with 10% FBS and 1X penicillin-streptomycin. Ovaries were washed three times in 1X PBS (Thermo Fisher: 70011044) and then fixed in 4% paraformaldehyde (Electron Microscopy Sciences: 15713) for 20 mins at RT. They were then washed three times in 1X PBS and stored in 100% methanol at 4℃ overnight or longer before smFISH experiments. Probe sequences for labeling *nos, pgc* and *gcl* mRNAs were described previously.^52^

#### smFISH in cells

After 24 hrs of transfection, cells on collagen-coated coverslips were fixed in 4% paraformaldehyde for 10 mins at RT and washed three times in 1X PBS. Coverslips were then stored in 70% ethanol (Thermo Fisher: BP2818500) at 4℃ overnight. The next day, samples were rehydrated in 1X PBS and washed three times in 1X PBS at RT. Cells were then permeabilized with 0.25% PBTx (1XPBS, 0.25% Triton-X100 (Millipore: TX1568-1)) for 3 mins at RT, followed by three washes in 1X PBS. Finally, samples were stained with smFISH probes with a modified protocol. Specifically, the probe hybridization was performed for 4-6 hrs at 37℃ instead of overnight before the washing step.

#### qRT-PCR for cells

After 24 hrs of transfection, cells were washed three times in chilled 1X PBS. Afterwards, 1 mL Trizol (Thermo Fisher: 15596026) was added directly to each well. The samples were then transferred to 1.5 mL microcentrifuge tubes and stored at −20℃ until use. For RNA extraction, 200 μL of chloroform was added to each tube and the RNA was purified using RNA Clean & Concentrator kit (Zymo: R1014) according to the manufacturer’s instructions. Approximately 1μg of total RNA was treated with RQ1 RNase-Free DNase (Promega: M6101) and cDNA was synthesized using SuperScript^TM^ III reverse transcriptase (Thermo Fisher: 18080093) following the manufacturer’s protocols. For each 10 μL qRT-PCR reaction, approximately 100 ng of cDNA, 1.5 μL of 1 μM forward and reverse primers, and 5 μL iTaq universal SYBR® Green supermix (Bio-Rad: 1725122) were combined. Quantitative PCR analysis was performed on a CFX Opus 96 Real-Time PCR System (Bio-Rad).

### QUANTIFICATION AND STATISTICAL ANALYSIS

#### Quantifying the number and intensity of condensates

Non-deconvolved images from vt-iSIM were used for condensate intensity analysis. For other experiments, images were deconvolved using Huygens Essential (Scientific Volume Imaging). The number and fluorescence intensity of condensates were quantified using Airlocalize spot detection algorithm as described previously.^23^

#### Western blot analysis

The analysis was performed according to the ImageJ User Guide (Section 30.13, Gels). For *Drosophila* samples and U2OS cells, protein levels were normalized to α-tubulin and β-actin loading controls, respectively.

#### FRAP analysis on Osk-eGFP granules in oocytes and embryos

The analysis was performed as described previously.^35^ Briefly, fluorescence recovery of ROIs was quantified in ImageJ. Recovery curves were generated using easyFRAP with full-scale normalization to account for differences in bleaching depth across experiments. Consequently, recovery curves were scaled from 0 to 1, with 1 corresponding to the pre-bleached ROI and 0 to the ROI immediately after photobleaching. As described previously,^35^ for live oocytes and embryos, the bleached ROI with the dimensions of 3×3 μm (please see above) was modeled as containing three populations: P1, a diffusive population residing in the intergranular space with fast mobility (M1); P2, a population located within granules that exchanges with the surrounding cytoplasm at a slower mobility (M2); and P3, the immobile fraction within granules that does not exchange. To quantify these populations, average recovery curves of normalized FRAP data were fitted to a two-term exponential equation (*f(t) = a^∗^(1-exp(-b^∗^t))+c^∗^(1-exp(d^∗^t))*), using Sigma plot (Grafiti). In this model, *a* and *c* represent the percent mobile fractions of P1 and P2, respectively, while *b* and *d* correspond to their recovery rate constants. From these constants, half-time recovery (*t_1/2_*) values for P1 and P2, denoted *t_½_* (P1) and *t_½_* (P2), were calculated using the equations *t_½_* (P1) = *ln(2)/b* and *t_½_* (P2) = *ln(2)/d*, respectively. FRAP data are included in Table S1.

#### FRAP analysis on Short Osk ΔNLS1-eGFP condensates in cells

The analysis was performed as described previously^26^ and the normalization was carried out as described above. Average recovery curves of normalized FRAP data were fitted to a single-term exponential equation (*f(t)=a*(1-exp(-b*t))*) using Sigma plot, where *a* represents the percent mobile fraction and *b* represents the recovery rate constant. *t_1/2_* was calculated using the equation *t_1/2_ = ln(2)/b*. FRAP data are included in Tables S2 and S3.

#### Co-localization analysis of germ granule mRNAs with Short Osk ΔNLS1-eGFP granules

Analyses were performed using PCC(Costes) using ImageJ plugin JACoP as described before.^35,41^

#### qRT- PCR analysis of germ granule mRNA levels in cells

Human *GAPDH* was used as a control to calculate the relative transcript levels, which was 2^−(Cq(gene of interest)− Average of Cq(*GAPDH*))^. Primers are listed in Table S4.

## ACKNOWLEDGMENTS

This research was supported by the NIGMS R35GM142737 awarded to T.T.

## Author contributions

Conceptualization: S.T. and T.T.; methodology: S.T., H.A.C. and T.T.; investigation and analysis: S.T., H.K. and H.A.C.; writing-original draft: S.T.; writing-review & editing: S.T. and T.T.; funding acquisition, resources and supervision: T.T..

## Declaration of generative AI and AI-assisted technologies in the writing process

During the preparation of this work, the authors use ChatGPT 5.1 to improve the readability and language of the manuscript.

